# Membrane constriction by dynamin through GTP-driven conformational changes from coarse-grained molecular dynamics simulations

**DOI:** 10.1101/2025.09.09.675253

**Authors:** Md. Iqbal Mahmood, Shintaroh Kubo, Hiroshi Noguchi, Kei-ichi Okazaki

**Affiliations:** Research Center for Computational Science, Institute for Molecular Science, National Institutes of Natural Sciences, Okazaki, 444-8585, Japan; Department of Cell Biology, Graduate School of Medicine, The University of Tokyo, Tokyo, Japan; Institute for Solid State Physics, The University of Tokyo, Kashiwa, Chiba 277-8581, Japan; Graduate Institute for Advanced Studies, SOKENDAI, Okazaki, Aichi 444-8585, Japan; Department of Physical Chemistry, School of Pharmacy and Pharmaceutical Sciences, Hoshi University

**Keywords:** Dynamin 1, Biomolecular machines, Membrane remodeling

## Abstract

Dynamin, a GTPase involved in membrane remodeling processes, plays a pivotal role in membrane fission during endocytosis. Understanding the molecular mechanisms of dynamin assembly and its conformational change triggered by GTP hydrolysis on lipid membranes is crucial for elucidating the intricate details of membrane constriction and fission events. However, the size and complexity of the dynamin assembly prevent us from obtaining comprehensive molecular mechanisms. In this study, we performed coarse-grained molecular dynamics (CG-MD) simulations to elucidate the chemo-mechanical coupling mechanism of dynamin membrane constriction. We set up a CG Martini simulation system that contains dynamin rings on a tubular membrane with realistic molecular structures. Controlling the length of the tubular membrane with a pressure coupling, we simulate membrane constriction through the nucleotide-state-dependent conformational change of dynamin. The simulation results suggest that the conformational change to the GDP state loosens and expands the dynamin rings, resulting in membrane constriction in the protein-uncoated region. This study lays the groundwork for simulating the chemo-mechanical coupling of dynamin-mediated membrane constriction.

**TOC GRAPHICS:** 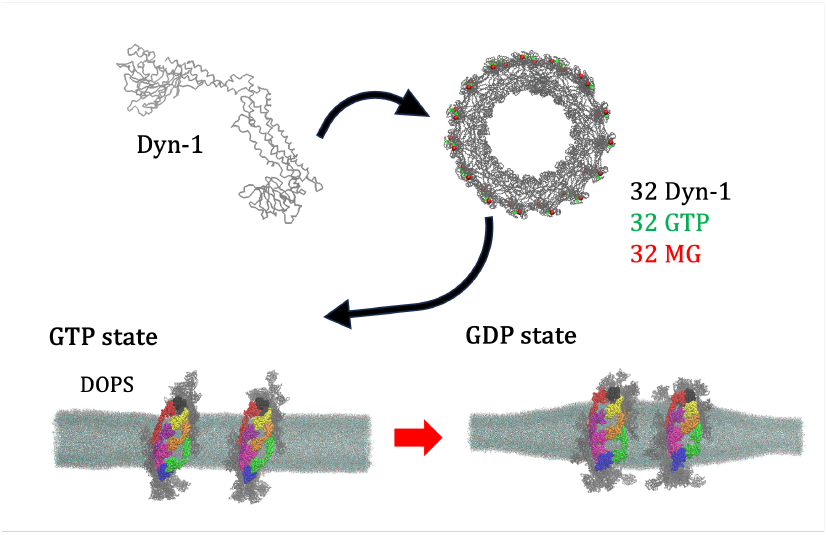

---

The large GTPase dynamin assembles on the neck of budding vesicles and catalyzes constriction and fission of the tubular membrane during endocytosis. X-ray crystallographic and cryo-electron microscopic (cryo-EM) studies have revealed structural insights into dynamin function^1,2^. Dynamin monomer has a GTPase (G) domain, a bundle signalling element (BSE), a stalk, a pleckstrin homology (PH) domain, and a proline-rich domain (PRD). The G-domain binds and hydrolyzes GTP, while the PH domain interacts with lipid membranes. Dynamin oligomerizes to form intricate helical assemblies, or lock-washer-like rings, on tubular membranes. Cryo-EM structures show three forms of helical assemblies: unconstricted, constricted, and super-constricted states^2–4^, which are observed in the apo, non-hydrolyzable GTP-analog, and GTP conditions, respectively. The super-constricted form features a two-start helix, where two helical chains of dynamin are interwound, resulting in an inner radius of the tube that is less than 2 nm. However, it is unclear whether the two-start helical assembly plays a role in dynamin-mediated fission of tubular membranes^4^.

Two different models have been debated in explaining dynamin-mediated membrane fission^3^. On one hand, the two-stage model assumes that constriction is mainly achieved through dynamin oligomerization, such as the constricted or super-constricted helical assembly. Then, the GTP-hydrolysis-driven conformational change destabilizes or loosens the assembly, inducing hemifission and subsequent fission of the tubular membrane. On the other hand, the constrictase (or ratchet) model assumes that the GTP-hydrolysis-driven conformational change is directly coupled to the mechanical work of constriction, which is analogous to the actomyosin system in the muscle^5^. The GTP-hydrolysis-triggered disassembly of dynamin oligomers is consistent with the two-stage model^3^, while kinetic measurements are consistent with the constrictase model^5^.

Due to the limitation in experimental spatial and temporal resolutions, molecular dynamics (MD) simulations have been employed to complement the experimental results^5,6^. In MD simulations, simplified models have been used rather than reproducing structural details of dynamin assemblies^5,7,8^. Pannuzzo *et al*. simplified the dynamin complex into a stack of beads, representing the controllable dynamin helical filament^8^. Ganichkin *et al*. simplified the dynamin filament into a chain of beads with explicit cross-bridges of the G-domain-G-domain (GG) interface^5^. Although these simplified models of the dynamin filament offer physical insights into the constriction mechanism of dynamin, they lack the structural aspects of individual dynamin monomers, potentially leading to unrealistic simulations of the dynamin filament.

To address this, we employ the coarse-grained (CG) Martini model, where each protein is represented by a coarse-grained structure with backbone and side-chain beads^9,10^. The CG MD simulations have been extensively applied to biological systems with lipid membranes and proteins^9,11^. It has also been applied to protein complex assembly^9,12–14^ and protein-induced membrane transformation^12,15–18^. In Martini, approximately four heavy atoms are coarse-grained into one bead, making it computationally more efficient than conventional all-atom MD simulations. Thus, the CG Martini approach is expected to facilitate the simulation of dynamin-induced membrane constriction.

In this study, we performed CG-MD simulations of the tubular membrane with the prototypical dynamin 1 (Dyn-1) rings (Figure 1A). The structure of the Dyn-1 ring was modeled from cryo-EM density, and LipidWrapper^19^ was used to build the tubular membrane (see SI text). With the CG-MD simulations, we investigate the dynamic behavior of the Dyn-1 complex and its interactions with the lipid membrane tubule. Then, we simulated the conformational change of Dyn-1 by switching the elastic network applied to each dynamin monomer. The simulation results provide insight into the intricate interplay among the conformational change of the Dyn-1 monomer, the Dyn-1 ring structure, and membrane deformation, advancing our understanding of cellular processes regulated by Dyn-1. In particular, our simulations determine if Dyn1 constricts the membrane directly or indirectly (Figure 1B).

**Figure 1.**
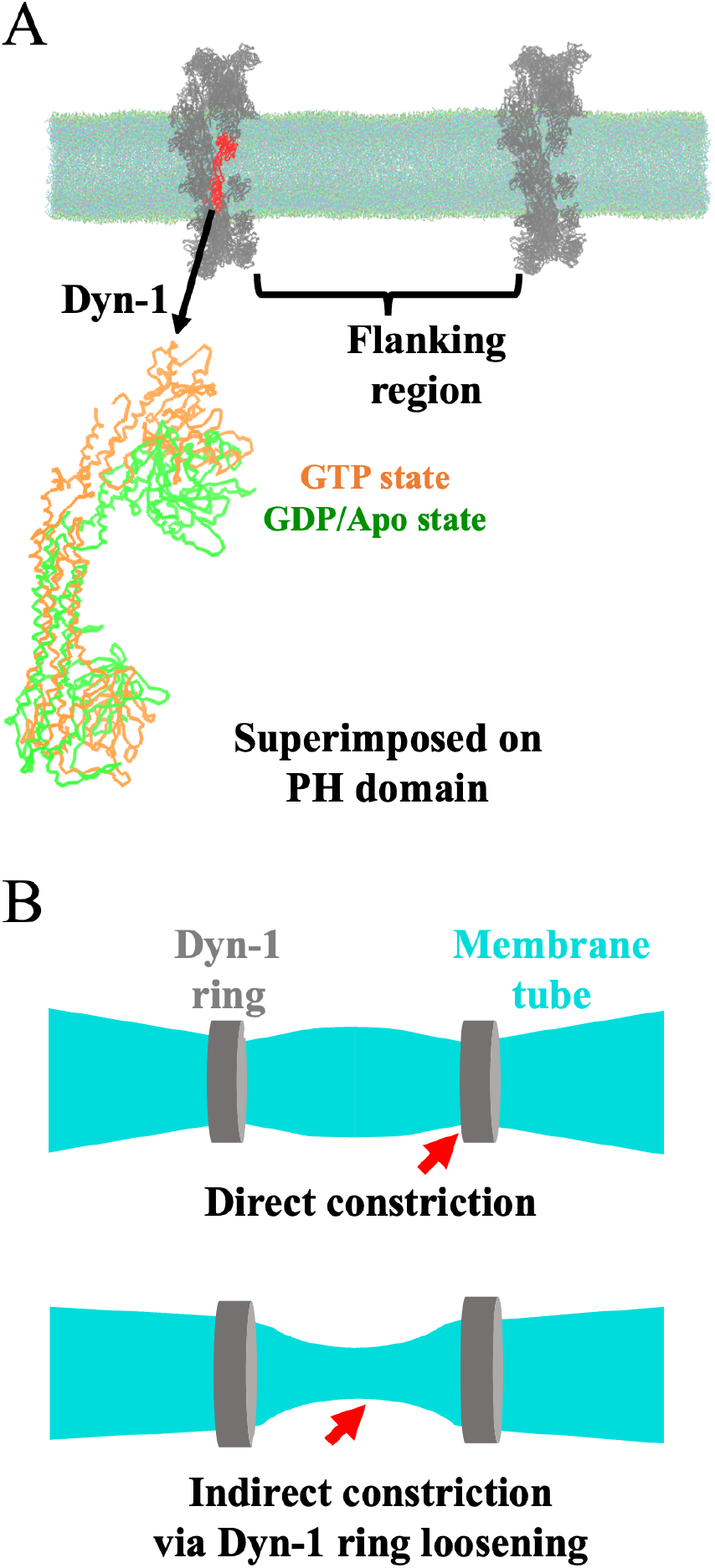
Dyn-1 assembly on tubular membrane and membrane constriction models. (A) The CG-MD simulation system of two Dyn-1 rings on a membrane tube is shown. With the PH domains aligned, the superimposed structures of the Dyn-1 monomer in the GTP-bound and GDP-bound/Apo states are depicted in orange and green, respectively. (B) Schematic representation of the Dyn-1 membrane constriction models. The direct constriction model, in which membrane constriction occurs at Dyn-1 ring positions, is shown at the top. The indirect constriction model, in which membrane constriction occurs at the flanking region, is shown at the bottom.

MD simulations of tubular membranes have rarely been performed^20,21^, and in some studies, they were performed without appropriate equilibration^22^. However, to simulate deformation of the tubular membrane, it is essential to control the tube length with a pressure coupling. Here, we simulated it with a semi-isotropic pressure coupling in which the long axis of the tube is controlled separately from the other two axes. During this simulation, we encountered the issue that the average pressure values significantly deviated from the target value of 1 bar (Figure S1). This fundamental issue turned out to be a general issue of simulating large systems with semi-isotropic or anisotropic pressure coupling, previously reported^23^. We addressed this problem by adopting the revised set of neighbor-list parameters in the MD engine GROMACS (see SI text). With the revised parameters, the pressure was controlled correctly (Figure 2).

**Figure 2.**
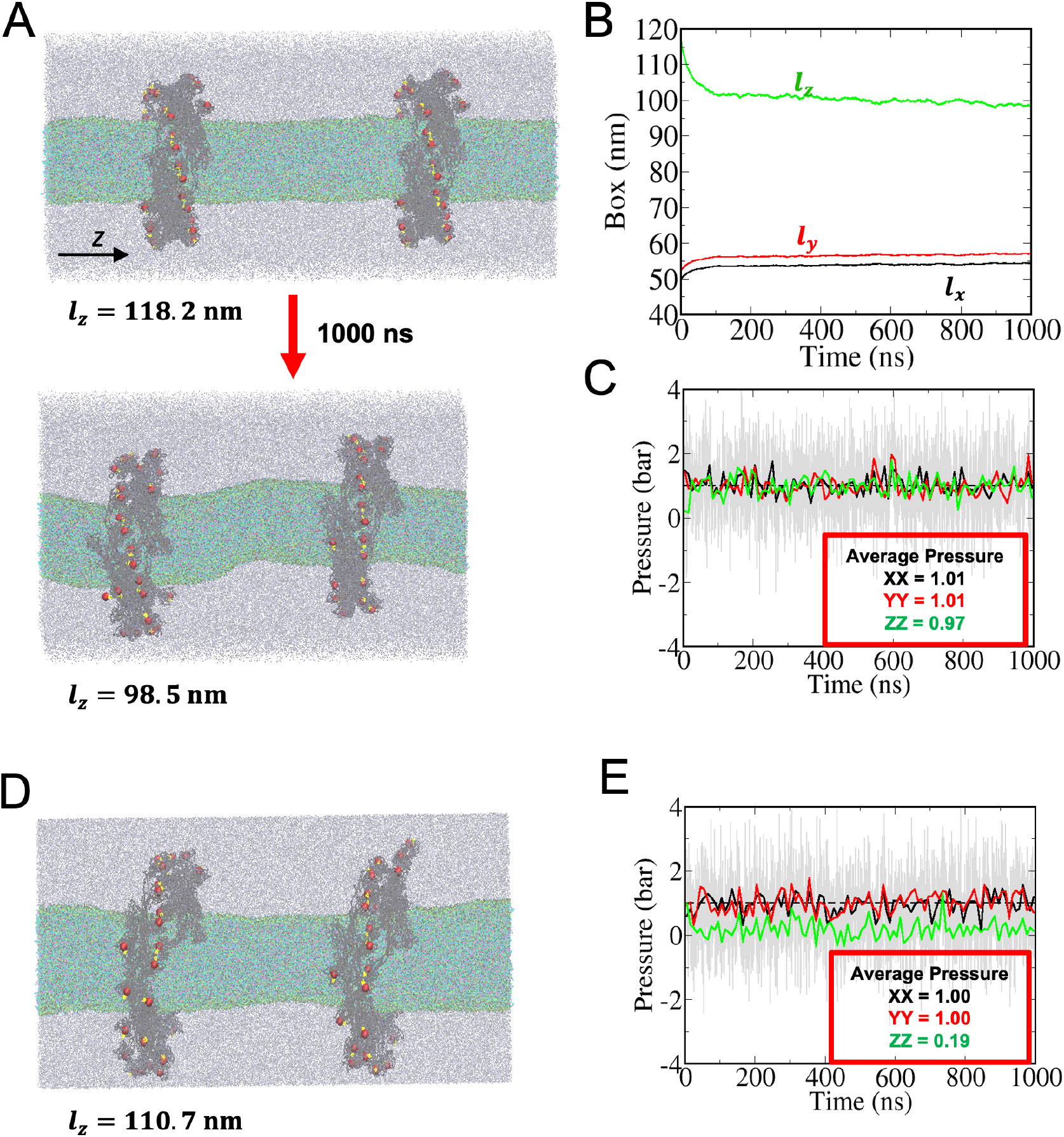
Corrected pressure control with the revised GROMACS parameters. (A) Initial and final configurations of the system at 1 bar. (B, C) Box lengths and pressures in the x, y, and z directions. (D) Final configuration of the system at 1 bar for the x and y directions and 0.2 bar for the z direction. (E) Pressures in the x, y, and z directions with z-axis pressure controlled at 0.2 bar instead of 1 bar.

Simulating with the corrected pressure control, we observed that the tubular membrane got bent with the standard pressure to the long axis of the tube (Figure 2A). When we reduced the target pressure of the long axis of the tube to 0.2 bar, it kept the straight tube shape (Figure 2D). This behavior can be understood based on the elastic theory of bilayer membranes^24–28^. For the energy of a tubular membrane *E*, we considered membrane bending elasticity, surface tension, area difference elasticity of the inner and outer leaflets, and work done by the pressure difference between the inside and outside the tube.

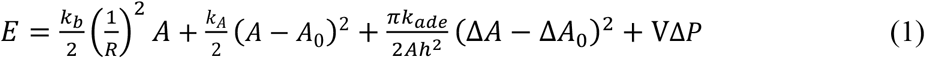

where *R* is the radius, *L* is the axial length, *V* = π*R*^2^*L* is the volume, *A* = 2π*RL* is the surface area of the tube, and Δ*P* = *P*_*out*_ − *P*_*in*_ is the pressure difference between outside and inside of the tube due to the water density difference. *k*_*b*_ is the constant for membrane bending elasticity, *k*_*ade*_ is the constant for area difference elasticity between the inner and outer leaflets^25,28^. *k*_*b*_ and *k*_*ade*_ are typically estimated as ∼20*k*_*B*_*T*^29,30^. *h* = 2 nm is the distance between the centers of the two leaflets, and Δ*A* = 2π*hL*, and Δ*A*_0_ is Δ*A* of the initial conformation. Using ∂E/ ∂R = 0 for adjusting the radius, the axial force is given (see SI text) by

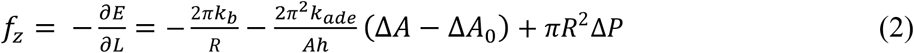

The axial force is converted to the pressure by dividing it by the xy area of the simulation box. First, the membrane bending elasticity contributes to the shrinkage of the tube, having a negative value of the axial force. The estimated pressure of this term, using the average tube radius of 7.5 nm from the reduced pressure simulation, was ∼ -0.26 bar. The contribution of the second area-difference elasticity of the inner and outer leaflets is small, with the estimated value of ∼ 0.06 bar. The third volume-pressure term contributes to the axial force when there is a density difference of water between the inside and outside the tube. The relative number density of water inside the membrane was ∼0.25 compared to actual water, which is close to the bulk value in Martini accounting for the four-to-one CG mapping (Figure S2)^31^. However, using the relation 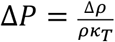 where *ρ* is the bulk water density, Δ*ρ* is the density difference between the inside and outside the tube, *k*_*T*_ is the compressibility of water, a small density difference leads to a large pressure difference. Thus, the contribution of this term would be significant in our case. We also note that the presence of Dyn-1 rings might also affect the axial force. Nonetheless, by lowering the target pressure in the axial direction, we can control the tube length and radius whose axial force accounts for the reduced target pressure.

Next, we simulate the conformational change of each Dyn-1 monomer from the GTP state to the GDP state, which has been considered a major driving force for membrane constriction^3,5^. Dyn-1 changes to a closed conformation in the GDP state, which is believed to generate a constriction force through the GG cross-bridge interaction. Here, we induced the conformational change by switching the elastic network (ELN) potential imposed on each Dyn-1 from the open to the closed conformation to see how it affects the conformation of the whole Dyn-1 ring and its interaction with the membrane. Similar approaches have been used to induce conformational changes in proteins^32,33^.

After the ELN switching, we observed conformational changes in the Dyn-1 ring conformation in the single Dyn-ring system. The four GG cross-bridges significantly changed the conformation at the ring edges (Figure 3A). The root mean square deviation (RMSD) plot clearly shows that the conformational change from the GTP to GDP state was completed (Figure 3C). At first sight, these changes seem consistent with the direct constriction by the Dyn-1 ring (Figure 1B). The PH domain positions after the ELN switching seem to suggest a tightening of the Dyn-1 ring (Figure 3B). However, the tubular membrane was not constricted at the Dyn-1 ring-coated position. Although the constriction was not significant for this tube length, it was slightly constricted in the region away from the Dyn-1 ring in one trajectory (see Figure S3). After a careful examination of the conformational change, we observed that the Dyn-1 filament was elongated at the ring edges. This can be quantified by the distance between two adjacent PH domains in the filament, which shows an increase for most of them (Figure 3D). Thus, the conformational change did not induce a direct constriction of the membrane.

**Figure 3.**
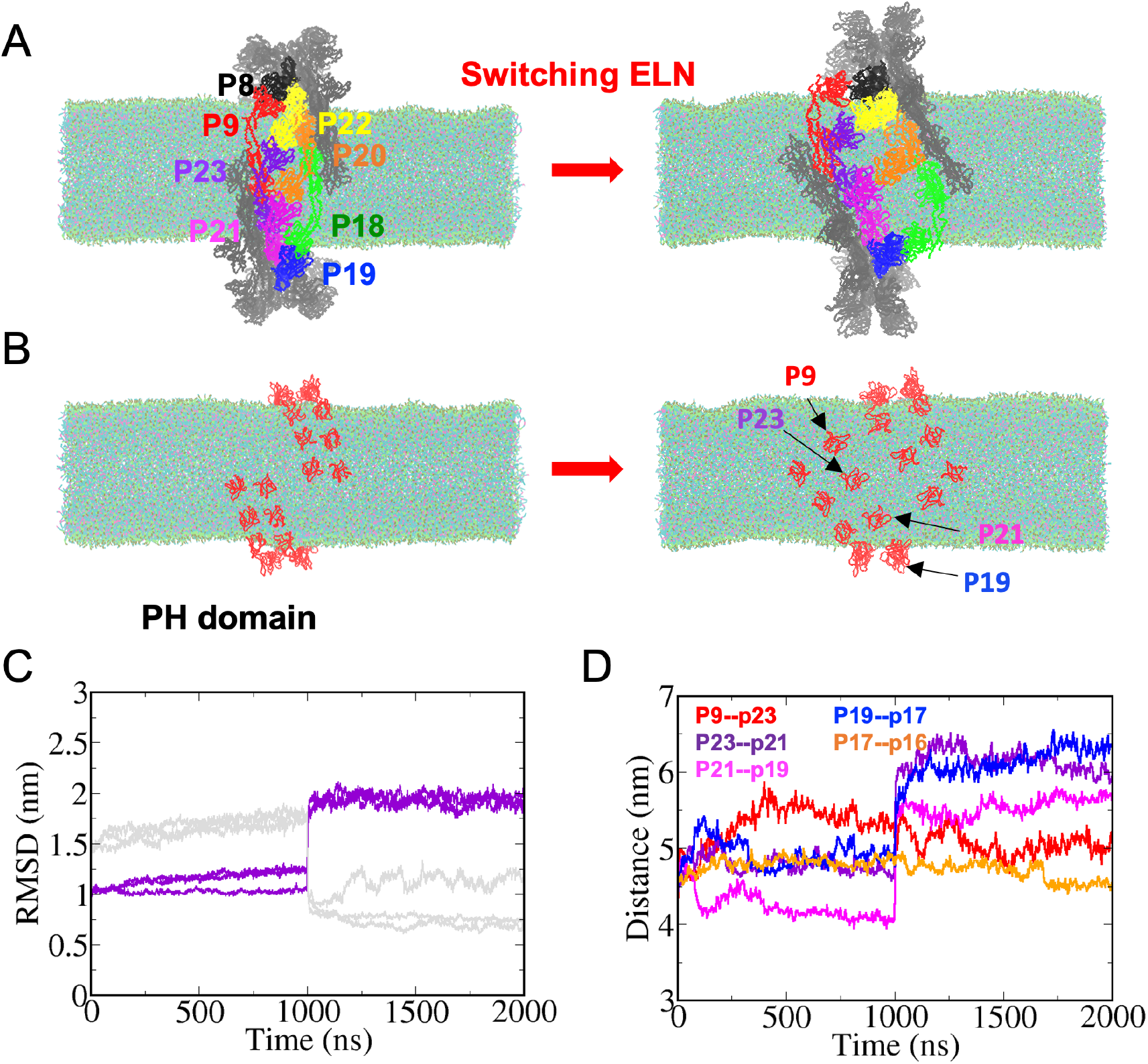
Conformational changes in Dyn-1 monomers induced by elastic network (ELN) switching. (A) Four GG cross-bridge interactions are highlighted, with each monomer represented in a distinct color. The Dyn-1 ring conformations before and after the ELN switching are shown. (B) The PH domain positions before and after the ELN switching are shown. (C) Time course of root mean square deviation (RMSD) for the monomer P23. The purple lines represent RMSDs to the open (GTP) conformation in three independent trajectories, while the gray lines represent RMSDs to the closed (GDP) conformation. The ELN was switched at 1000 ns. (D) The distances between PH domains located at the ring edge are shown.

Then, we simulated the conformational changes in the double Dyn-1 ring system. After the conformational changes induced by the ELN switching, the membrane tube was constricted in the protein-uncoated flanking region (Figure 4A). The membrane tube radius, estimated by fitting lipid beads, significantly decreased by 2 to 3 nm at the protein-uncoated region (Figure 4B and 4C). Thus, the conformational changes of the double Dyn-1 rings induced an indirect membrane constriction, which is consistent with the single Dyn-1 ring system. The membrane constriction is more prominent in the double Dyn-1 ring system. The mechanism of how the Dyn-1 ring induces membrane constriction at the uncoated region can be explained by the increase in the radius of gyration of each ring after the conformational change (Figure 4D). This expansion of the Dyn-1 ring induces an inflation of the membrane tube at the Dyn-1-coated region (Figure 4B and 4C), leading to constriction at the uncoated flanking region. The constriction at the dynamin-uncoated flanking region is consistent with the previous high-speed atomic force microscopy (HS-AFM) experiment^34^.

**Figure 4.**
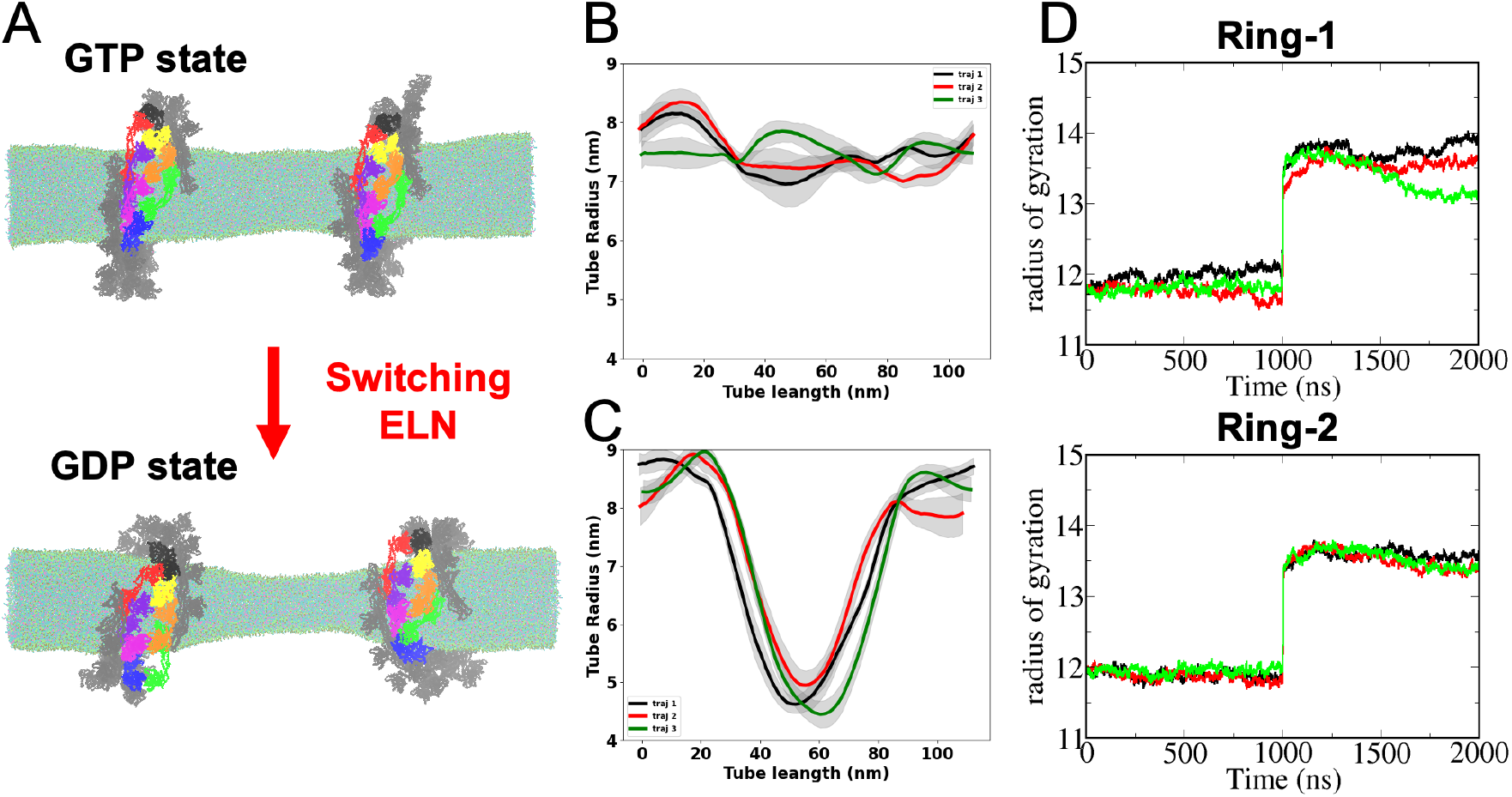
Tubular membrane constriction by conformational changes of Dyn-1 rings. (A) Snapshots of the GTP and GDP states are shown, highlighting structural differences between the two forms. (B) Average membrane tube radius of the GTP state along the tube long axis during three independent trajectories. (C) Average membrane tube radius of the GDP state along the tube long axis during three independent trajectories. The tube radius was calculated by fitting a circle to the xy positions of the PO4, C4A, and C4B beads of the DOPS lipid molecules for the last 500ns of the MD simulations. (D) The radius of gyration for two Dyn-1 rings from the GTP state to the GDP state over the course of three independent MD simulations is plotted.

In Figure 4A, the tubular membrane was constricted at the center, not at the edges. We investigated this apparent asymmetry by changing the relative position of the two Dyn-1 rings. Assuming that the tubular membrane is constricted at a wider uncoated region, we set up a system where two Dyn-1 rings are closer at the center. In this case, the tubular membrane was indeed constricted at the edges (Figure 5). This result justifies our assumption that the tubular membrane is constricted at a wider uncoated region. Based on our results, we propose a mechanism in which the expansion of the Dyn-1 rings induces different tube radii in the coated and uncoated regions, leading to the formation of a cluster of Dyn-1 complexes (Figure 6). We also simulated a system with a single Dyn-1 ring with double helical turns at the center to obtain similar results (Figure S4).

**Figure 5.**
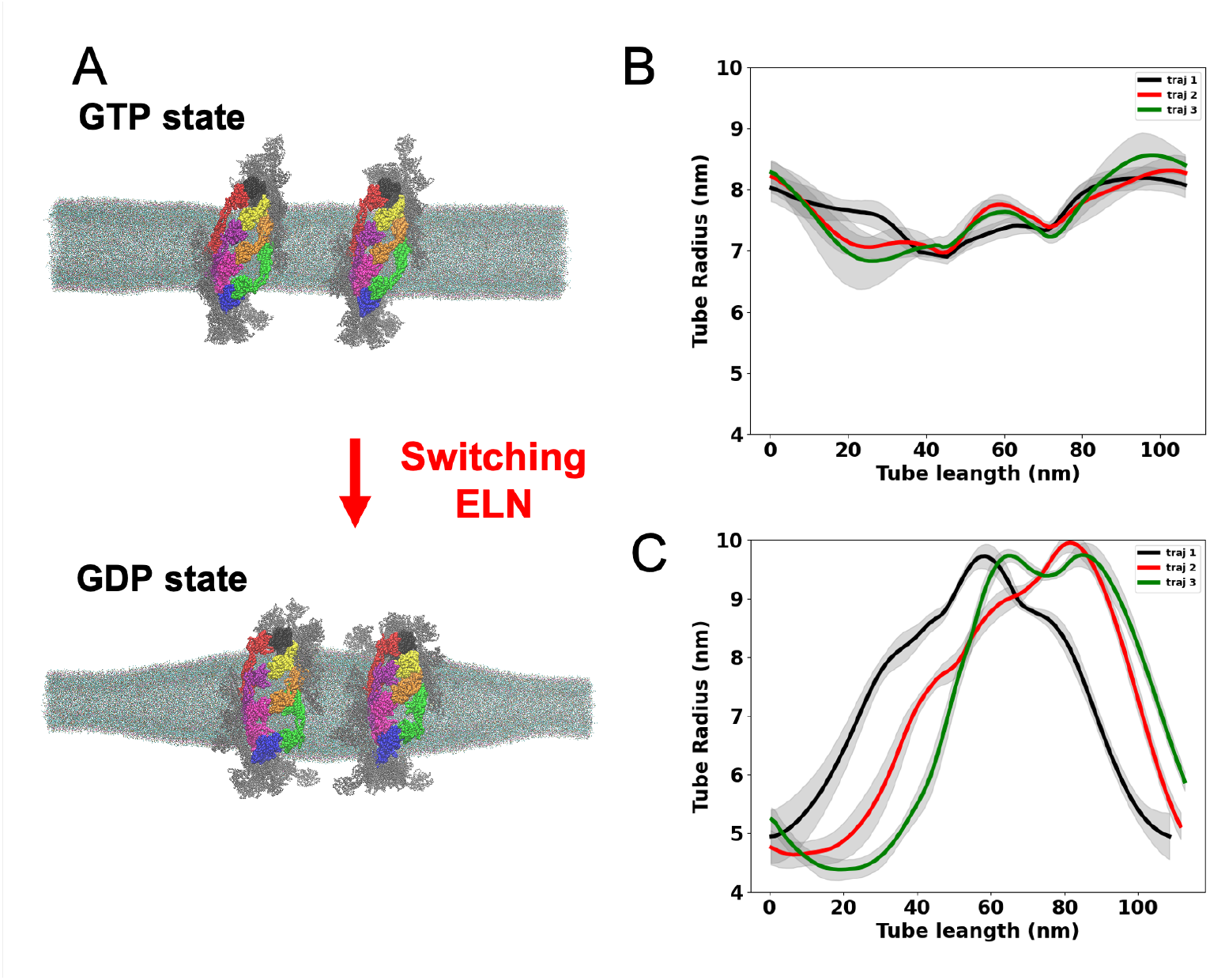
Tubular membrane constriction by conformational changes of Dyn-1 rings closely placed at the center. (A) Snapshots of the GTP and GDP states are shown, highlighting structural differences between the two forms. (B) Average membrane tube radius of the GTP state along the tube long axis during three independent trajectories. (C) Average membrane tube radius of the GDP state along the tube long axis during three independent trajectories. The tube radius was calculated by fitting a circle to the xy positions of the PO4, C4A, and C4B beads of the DOPS lipid molecules for the last 500ns of the MD simulations.

**Figure 6.**
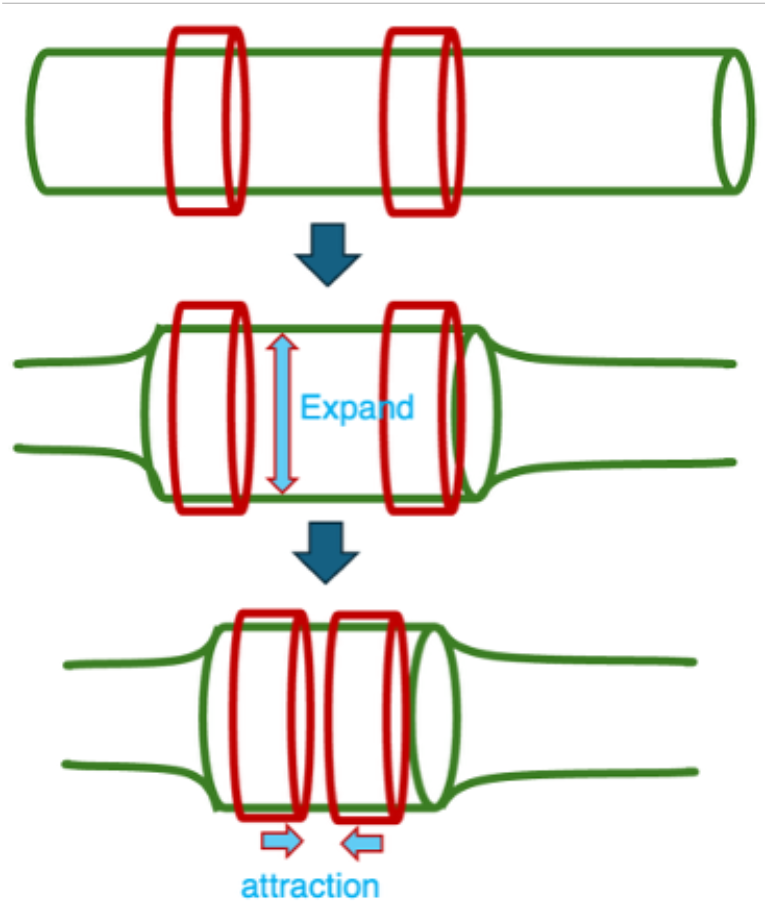
Mechanism of tubular membrane constriction by Dyn-1. The red and green objects represent the Dyn-1 ring and the tubular membrane, respectively.

In summary, we demonstrate that the conformational change of Dyn-1 monomers upon GTP hydrolysis results in an indirect constriction of the tubular membrane, as observed through CG-MD simulations. Contrary to the common notion that the Dyn-1 ring directly constricts the membrane by tightening the ring itself, our simulations suggest that the Dyn-1 ring loosens and expands upon GTP hydrolysis to constrict the membrane at the protein-uncoated region (Figure 6). Although our simulations provide insight into the constriction mechanism of dynamin, we have not yet simulated the complete fission of the tubular membrane. There are several challenges to simulating the complete fission. In our current simulations, the tubular membrane with its long axis connected through the periodic boundary condition is filled with water, and its pressure acts against the fission. Simulating a large vesicle with a tubular neck may resolve this problem^8^, although this requires a larger simulation system with higher computational costs. Another challenge involves the conformational change of Dyn-1 monomers. With the concerted transition to the GDP state assumed in the current simulations, conformational intermediates are not considered. Modeling intermediate conformations from HS-AFM experiments^34,35^ would help simulate realistic conformational changes. Nonetheless, our CG-MD simulations with a realistic Dyn-1 structure pave the way for understanding the chemo-mechanical coupling mechanism of Dyn-1 membrane constriction.

## ASSOCIATED CONTENT

### Supporting Information

Additional computational and methodological details; Deviation of average pressure from the target with the default Gromacs parameters (Figure S1); Water density across the XY plane (Figure S2); Tube radius of the GTP and GDP states (Figure S3); Tubular membrane constriction by conformational changes of the Dyn-1 ring with double helical turns placed at the center (Figure S4); Lipid tube preparation process (Figure S5) (PDF)

Simulation movie of membrane constriction by Dyn-1 conformational changes (MPG)

## AUTHOR INFORMATION

### Corresponding Author

* Kei-ichi Okazaki – Research Center for Computational Science, Institute for Molecular Science, National Institutes of National Sciences, Okazaki 444-8585, Japan; Graduate Institute for Advanced Studies, SOKENDAI, Okazaki, Aichi 444-8585, Japan;

### Notes

The authors declare no competing financial interest.

## ACKNOWLEDGMENT

The computation was partly performed using the Research Center for Computational Science, Okazaki, Japan (Project: 22-IMS-C189, 23-IMS-C201, 24-IMS-C198, and 25-IMS-C227). We thank Drs. A. Schahl and P. C. T. Souza for providing us with GTP and MG models. K.O. thanks Drs. R. Iino and J. Ohnuki for the helpful comments on this work.

## ABBREVIATIONS

Dyn-1: dynamin 1
CG-MD simulation: coarse-grained molecular dynamics simulation.

