## Supporting information for "Membrane constriction by dynamin through GTP-driven conformational changes from coarse-grained molecular dynamics simulations"

AUTHOR INFORMATION

**Corresponding Author**

*

### Computational Methods

*Cryo-EM modeling of Dynamin-1 assembly*

Modeling of Dyn-1 was performed based on PDB: 6DLU[1]. For the modeling of 32-mer Dyn-1, 32 copies of the structure were prepared and fitted into the density map (EMD: 7957)[1]. Each copy was placed within the map using the *fitmap* function in UCSF ChimeraX[2]. The final models were then adjusted to the density using Coot[3] and refined in real space in Phenix[4].

*CG-MD system setup of Dynamin-1 with tubular membrane*

We performed CG-MD simulations employing the latest version of the MARTINI force field Martini3[5]. We first prepared a flat lipid bilayer using the "insane.py" script[6]. The bilayer was composed of DOPS lipids, with 100% DOPS content, as this composition has been shown to promote the formation of extended dynamin-coated tubules, as observed in electron microscopy studies[7]. To create a tubular membrane structure from the flat bilayer, we employed the "lipidwrapper.py" script[8]. To avoid crashes of lipids when generating the entire membrane tube at once, the upper and lower halves, sliced along the tube axis, were generated separately and combined after equilibration (Figure S5). The membrane-tube inner diameter was ~10 nm.

The starting structure of the Dyn-1 protein for the MD simulation was obtained from the Cryo-EM structure of the membrane-associated helical filament of GTP-bound human Dyn-1 (PDB ID: 6DLU)[1]. The complete assembly was modeled from the cryo-EM density data as described above. The crystal structure of nucleotide-free human Dyn-1 (PDB ID: 3SNH)[9] was used for the GDP/apo reference structure for the ELN switching. Missing residues in the Dyn-1 GTP-bound dimer structure (PDB ID: 6DLU) were modeled using MODELLER[10,11]. The quality of the modeled structure was evaluated using the discrete optimized protein energy (DOPE)[12] score provided by MODELLER. Specifically, missing residues for the first monomer included M1-M6 and S21-D28, while the second monomer was missing T748. Similarly, missing residues in the nucleotide-free Dyn-1 structure (PDB ID: 3SNH) were also modeled with MODELLER, which included M1-G5, I63, V64, G110-N112, V143-P149, S347-L356, N394-T404, L446, Q447, Q500-E517, M534-G537, G578-S581, D632-D652, and G709-M715. We then generated the dynamin assembly ring with 32 Dyn-1 monomers, having four GG cross-bridges. After generating the dynamin assembly ring, the all-atom model was converted to a coarse-grained (CG) MARTINI model using the Martini force field v3.027, following the parameters and protocols established by the MARTINI community, with the aid of an elastic network. The "martinize2"[13] was employed to generate the coarse-grained structure and topology of the protein. A GTP molecule and a magnesium ion ($Mg^{2+}$) were added to each G-domain of Dyn-1. Distance restraints were applied between the beta phosphate of GTP and $Mg^{2+}$ to prevent $Mg^{2+}$ dissociation from the G-domain. We then placed the Dyn-1 assembly ring onto the membrane tube. The system was subsequently solvated with coarse-grained water molecules and ionized with 150 mM NaCl employing the "insane.py" script[6]. We designed two systems for our study. System 1 consists of a 60 nm-long membrane tube with a single dynamin assembly ring containing 32 dynamin monomers, resulting in a total of 1,284,994 beads. System 2 consists of a 120 nm-long membrane tube with two separate dynamin assembly rings, each comprising 32 dynamin monomers, totaling 2,678,021 beads.

*MD simulations*

In this study, we used the MD engine GROMACS 2021 (www.gromacs.org)[14,15]. To address the problem of a significant deviation of the average pressure during MD simulations under the default GROMACS parameters, we revised the parameters based on the recommendations by Kim et al.[16] as follows: *nstlist* was set to 20, and the *Verlet-buffer-tolerance* (VBT) was adjusted to approximately 0.0002 kJ·mol$^{-1}$·ps$^{-1}$. These adjustments ensured system stability during the simulations. The systems were first energy-minimized, followed by equilibration in the NPT ensemble with the semi-isotropic pressure coupling. Subsequently, production simulations were carried out for a duration of 1 μs. The system was maintained at 310 K using a v-rescale thermostat[17]. Pressure coupling was applied semi-isotropically with a C-rescale barostat[18], targeting 1 bar for the x-y axes and 0.2 bar for the z-axis. To investigate structural changes, the Martini elastic network topology was switched from the GTP-bound state (PDB ID: 6DLU) to the GDP-bound state (PDB ID: 3SNH), followed by 1 μs MD simulations.

*Axial force from the elastic theory*

The axial force in the main text was derived from the following equations.

$$\frac{\partial E}{\partial R} = -\frac{\pi k_b L}{R^2} + 2\pi\, L\sigma - \frac{\pi k_{ade}}{2Ah^2 R}(\Delta A - \Delta A_0)^2 + 2\pi R L \Delta P = 0 \qquad \text{(S1)}$$

$$f_z = -\frac{\partial F}{\partial L} = -\frac{\pi k_b}{R} - 2\pi\, R\sigma + \frac{\pi k_{ade}}{2Ah^2 L}(\Delta A - \Delta A_0)^2 - \frac{2\pi^2 k_{ade}}{Ah}(\Delta A - \Delta A_0) - \pi R^2 \Delta P \qquad \text{(S2)}$$

where surface tension $\sigma = k_A(A - A_0)$.

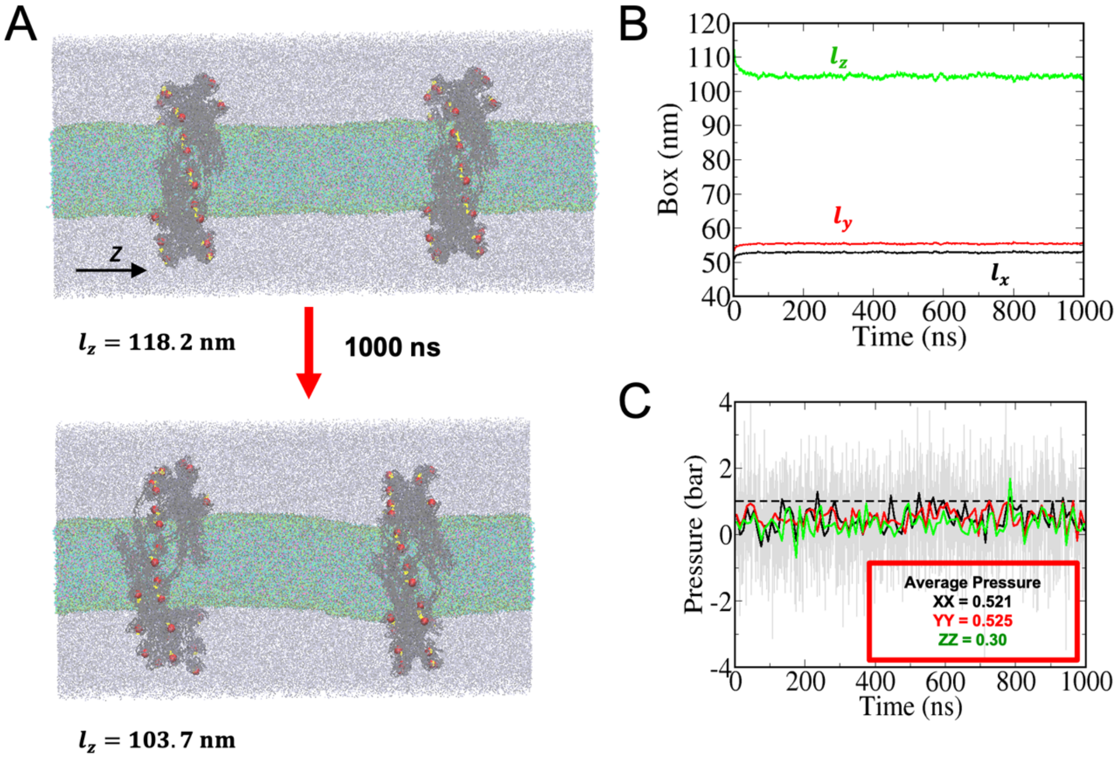


**Figure S1.** Deviation of average pressure from the target with the default Gromacs parameters. (A) The initial and final configurations of the system. (B) Evolution of the box lengths during the MD simulations. (C) Average pressure in the x, y, and z directions throughout the MD simulations. The solid lines represent running averages. The broken line represents the target pressure.

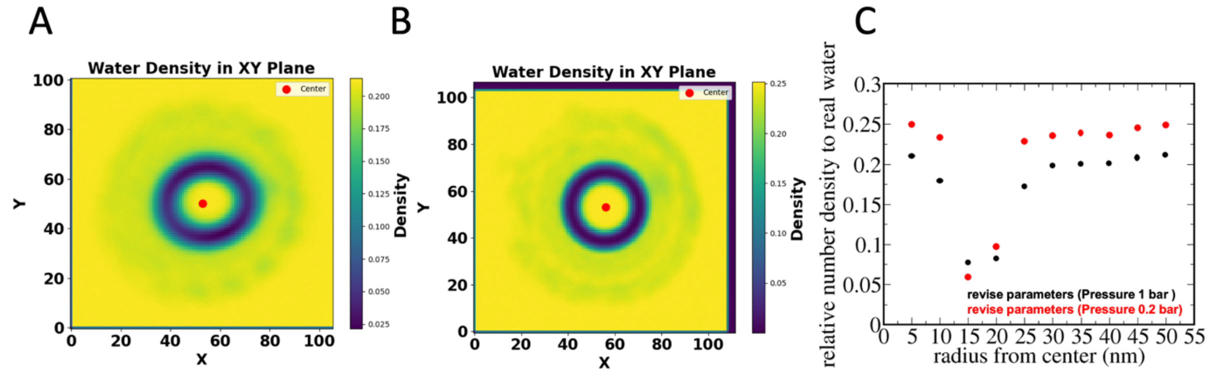


**Figure S2.** Water density across the XY plane. a) Water density distribution for the system using revised GROMACS parameters with Z-axis pressure controlled at 1 bar. b) Water density distribution for the system using revised GROMACS parameters with Z-axis pressure controlled at 0.2 bar instead of 1 bar. c) Comparison of water density between the two revised GROMACS parameter systems, represented by black and red colors, respectively.

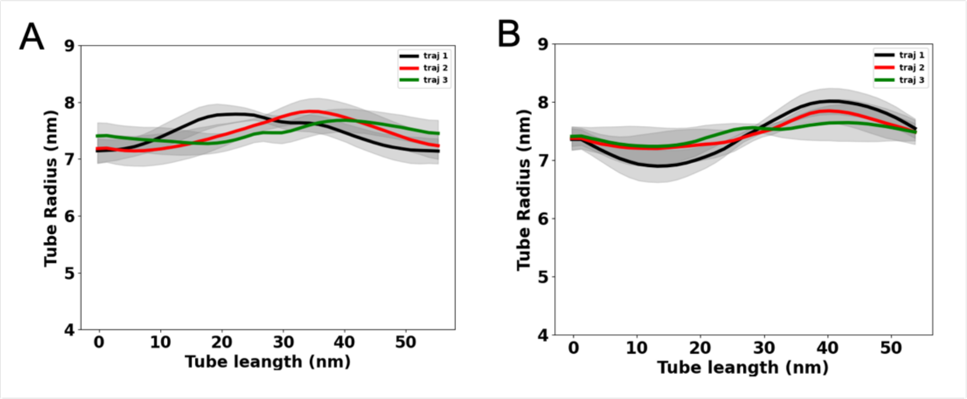


**Figure S3.** Tube radius of the GTP and GDP states. A) The tube radius of the GTP state along the tube axis during the three MD trajectories is shown in different colors. B) The tube radius of the GDP state along the tube axis during the three MD trajectories is shown in different colors. The radius is determined by fitting a circle to the positions of the PO4, C4A, and C4B beads of the lipid molecules.

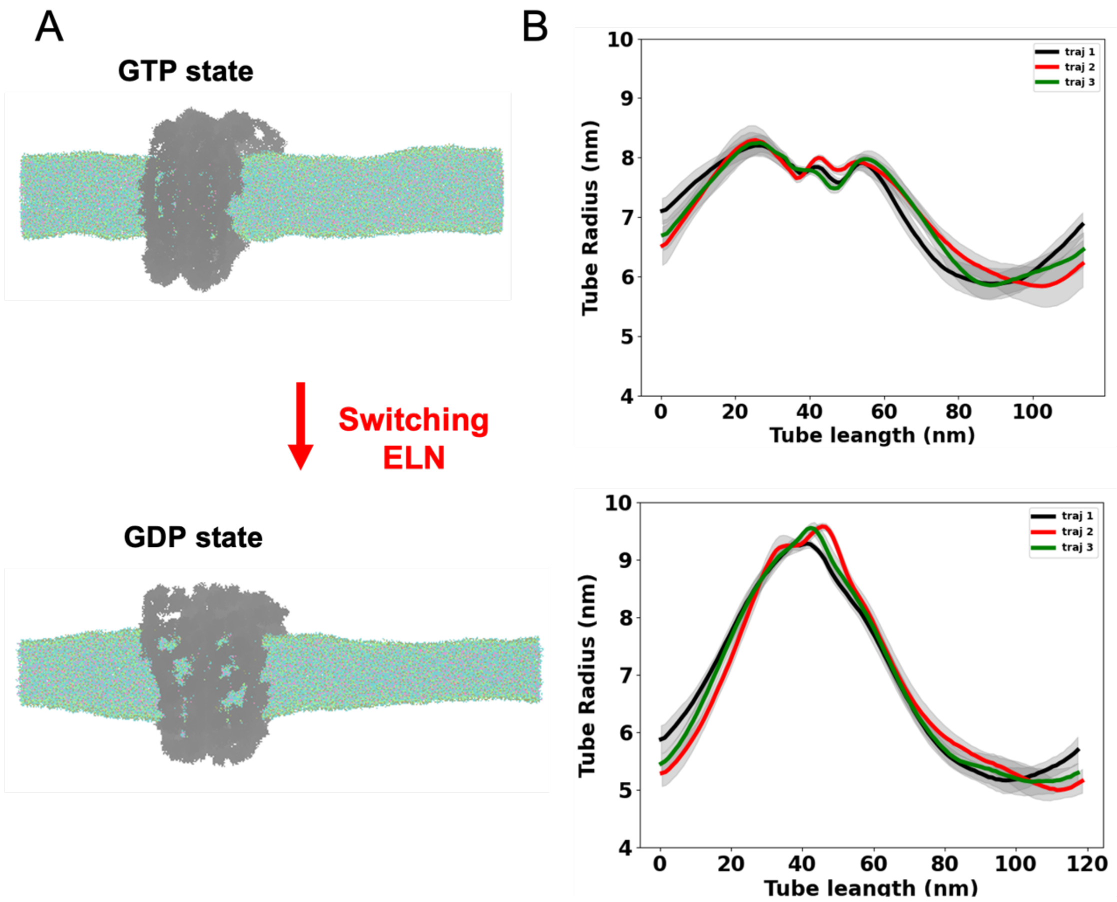


**Figure S4.** Tubular membrane constriction by conformational changes of the Dyn-1 ring with double helical turns placed at the center. (A) Snapshots of the GTP and GDP states are shown, highlighting structural differences between the two forms. (B) Average membrane tube radius of the GTP state along the tube long axis during three independent trajectories. (C) Average membrane tube radius of the GDP state along the tube long axis during three independent trajectories. The tube radius was calculated by fitting a circle to the xy positions of the PO4, C4A, and C4B beads of the DOPS lipid molecules for the last 500ns of the MD simulations.

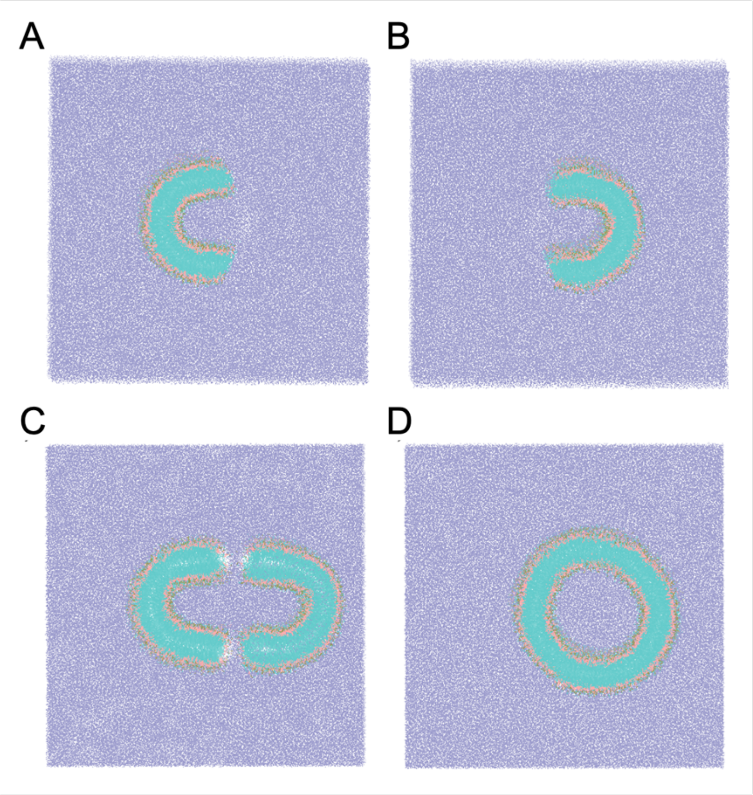


**Figure S5.** Lipid tube preparation process. The upper half (A) and lower half (B) tubes are shown. They were equilibrated for 0.5 ns separately. C) Both halves are merged and placed in a solvent environment. D) The lipid layers self-assembled into a tubular structure during the 100 ns equilibration process. The lipidwrapper.py script generated tube halves.
